# Establishment of regions of genomic activity during the *Drosophila* maternal to zygotic transition

**DOI:** 10.1101/006312

**Authors:** Xiao-Yong Li, Melissa M. Harrison, Tommy Kaplan, Michael B. Eisen

**Author notes:** co-senior authors.

## Abstract

A conspicuous feature of early animal development is the lack of transcription from the embryonic genome, and it typically takes several hours to several days (depending on the species) until widespread transcription of the embryonic genome begins. Although this transition is ubiquitous, relatively little is known about how the shift from a transcriptionally quiescent to transcriptionally active genome is controlled. We describe here the genome-wide distributions and temporal dynamics of nucleosomes and post-translational histone modifications through the maternal-to-zygotic transition in embryos of the pomace fly *Drosophila melanogaster*. At mitotic cycle 8, when few zygotic genes are being transcribed, embryonic chromatin is in a relatively simple state: there are few nucleosome free regions, undetectable levels of the histone methylation marks characteristic of mature chromatin, and low levels of histone acetylation at a relatively small number of loci. Histone acetylation increases by cycle 11, but it is not until cycle 13 that nucleosome free regions and domains of histone methylation become widespread. Early histone acetylation is strongly associated with regions that we have previously shown are bound in early embryos by the maternally deposited transcription factor Zelda. Most of these Zelda-bound regions are destined to be enhancers or promoters active during mitotic cycle 14, and our data demonstrate that they are biochemically distinct long before they become active, raising the possibility that Zelda triggers a cascade of events, including the accumulation of specific histone modifications, that plays a role in the subsequent activation of these sequences. Many of these Zelda-associated active regions occur in larger domains that we find strongly enriched for histone marks characteristic of Polycomb-mediated repression, suggesting a dynamic balance between Zelda activation and Polycomb repression. Collectively, these data paint a complex picture of a genome in transition from a quiescent to an active state, and highlight the role of Zelda in mediating this transition.

## Introduction

In most animals, the first phase of embryonic development depends solely on maternally deposited proteins and RNAs and is often accompanied by very low or undetectable transcription (Newport and Kirschner, 1982a, b; Tadros and Lipshitz, 2009). After several hours to several days, depending on the species, zygotic transcription initiates, marking the beginning of a process known as the maternal-to-zygotic transition (MZT) during which maternally deposited RNAs are degraded and the zygotic genome assumes control of its own mRNA production.

In *Drosophila melanogaster*, sustained zygotic transcription begins around mitotic cycle 7, about an hour into development, although there is growing evidence that very low levels of transcription occur even earlier (Ali-Murthy et al., 2013; ten Bosch et al., 2006). Zygotic transcription gradually increases with each subsequent mitotic cycle, but it is not until the end of mitotic cycle 13 that widespread zygotic transcription is observed (Lécuyer et al., 2007; Lott et al., 2011; McKnight and Miller, 1976; Pritchard and Schubiger, 1996). This zygotic genome activation, along with the elongation of mitotic cycle, and cellularization of the syncytial nuclei defines the mid-blastula transition (MBT). Approximately 3,000 genes are transcribed in the cellular blastoderm (De Renzis et al., 2007; Lécuyer et al., 2007; Lott et al., 2011). Of these, roughly 1,000 are expressed in spatially restricted patterns (Combs and Eisen, 2013; Tomancak et al., 2007), a product of the binding and activity of around fifty spatially restricted transcription factors to several thousand known and putative patterning enhancers (Li et al., 2008; MacArthur et al., 2009).

In the cellular blastoderm, active genomic regions are biochemically distinct from the rest of the genome: they have relatively low nucleosome densities; are bound by transcription factors, polymerases and other proteins that mediate their activity; and have characteristic histone modifications (Li et al., 2011; Nègre et al., 2011). This high level of activity and relatively complex landscape of genome organization is remarkable given that an hour earlier the genome was being continuously replicated and doing little else. Although the *Drosophila* cellular blastoderm is among the most well-characterized animal tissues, the transition from quiescent to active state that precedes the formation of this tissue remains poorly understood, despite increasing evidence of its importance (Blythe et al., 2010; Harrison et al., 2011; Lee et al., 2013; Leichsenring et al., 2013; Liang et al., 2008; Liang et al., 2012; Nien et al., 2011).

We have previously shown that a single seven base-pair sequence is found in the vast majority of patterning enhancers active in the cellular blastoderm (Li et al., 2008), and that the maternally deposited transcription factor Zelda (ZLD) (Liang et al., 2008), which binds to this sequence, is present at these enhancers by mitotic cycle 8, in the early phase of the MZT (Harrison et al., 2011). ZLD and its binding site are also found in the promoters of most genes activated during the MZT (ten Bosch et al., 2006), suggesting that it may play a broad role in early embryonic genome activation, analogous to pioneer transcription factors that choreograph the reorganization of genome activity during differentiation [reviewed in (Zaret and Carroll, 2011).

Although ZLD mutants alter the expression of a large number of cellular blastoderm genes (Liang et al., 2008), and affect transcription factor binding in the cellular blastoderm (Xu et al., 2014), little is known about its molecular function or when its activity is required. To gain further insights into ZLD’s mechanism and to place its action in the broader context of the MZT, we have broadly characterized the chromatin landscape throughout the *Drosophila melanogaster* MZT.

## Results

### Quantitative mapping of nucleosome occupancy and histone modifications across the MZT

To define the chromatin landscape before, during and after the maternal-to-zygotic transition, we collected *D. melanogaster* (Oregon-R) embryos from population cages at 25°C for 30 minutes, and aged them for 55, 85, 120 and 160 minutes to target mitotic cycles 8, 11, 13 and 14 respectively prior to fixing them with formaldehyde.

As *D. melanogaster* females often retain eggs post-fertilization, leading to unacceptable levels of contaminating older embryos in embryo pools (Harrison et al., 2011), we manually removed embryos of the incorrect stage by inspection under a light microscope (Figure 1A), as previously described (Harrison et al., 2011). The purity of the resulting embryo pools was confirmed by examining the density of nuclei in 4′,6-diamidino-2-phenylindole (DAPI) stained samples from each pool (Figures 1B and 1C).

**Figure 1.**
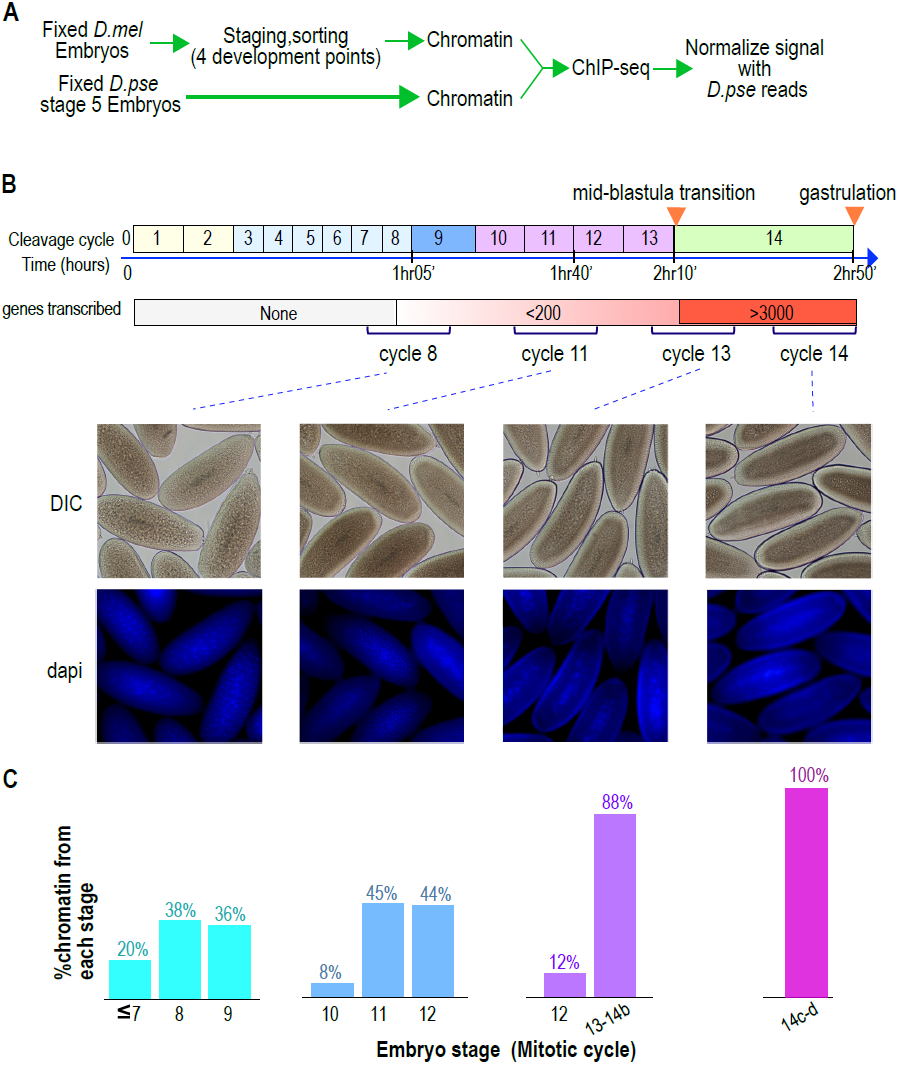
Hand sorting based on morphology results in tightly staged embryos. (A) Experimental scheme. *D. melanogaster* embryos were collected and allowed to develop before being fixed with formaldehyde. Fixed embryos were hand sorted to obtain pools of embryos within a relatively narrow age distributions between mitotic cycle 8 and the end of cycle 14. To serve as carrier and normalization standard, chromatin from fixed stage 5 (cycle 14) *D. pseudoobscura* embryos was prepared and added to the chromatin from the sorted embryos prior to chromatin immunoprecipitation. In ChIP-seq data analysis, the sequencing reads for *D. pseudoobscura* were used to normalize the *D. melanogaster* ChIP-seq signals. (B) Embryo collection and sorting. The timeline of the early embryogenesis is depicted on top with the relative lengths of each mitotic cycle approximated by the size of the box. The developmental stages (from 1-5) are indicated by different colors. The earliest sustained transcription is detected is at cycle 7, and the mid-blastula transition (MBT) occurs when a large number of genes are transcriptionally activated at approximately the end of cycle 13. We generated four pools of sorted embryos with developmental stages centered around cycles 8, 11, 13, or 14 as shown by differential interference contrast (DIC) and DAPI. (C) We determined the distribution of the developmental cycle of the embryos in each pool as shown by counting the number of nuclei in DAPI-stained embryos or by examining the extend of membrane envagination during cycle 14.

We carried out chromatin immunoprecipitation and DNA sequencing (ChIP-seq) using commercial antibodies against nine post-translation modifications (acetylation at H3K9, H3K18, H3K27, H4K5 and H4K8, mono-methylation at H3K4, and tri-methylation at H3K4, H3K27 and K3K36), as well as histone H3 (Table 1).

**Table 1.**

| ***Mark*** | ***AB source*** | ***AB catalog #*** |
| --- | --- | --- |
| H3 | Abcam | ab1791 |
| H4K5ac | Millipore | 07-327 |
| H4K8ac | Abcam | ab15823 |
| H3K18ac | Abcam | ab1191 |
| H3K27ac | Abcam | ab4729 |
| H3K4me1 | Abcam | ab8895 |
| H3K4me3 | Abcam | ab8580 |
| H3K9ac | ActiveMotif | 39138 |
| H3K27me3 | Millipore | 07-449 |
| H3K36me3 | Abcam | ab9050 |

As we sought to compare not just the genomic distribution of marks but also their relative levels across the MZT, we prepared chromatin from stage 5 *D. pseudoobscura* embryos (mitotic cycle 14), and added a fixed amount to each *D. melanogaster* chromatin sample prior to ChIP to serve as both a quality control and as a normalization standard. Since, for a given antibody, we expect the *D. pseudoobscura* chromatin in each time point to be identically immunoprecipitated (within experimental error), we could both evaluate the success of the immunoprecipitation, and compute the relative abundance of each histone modification over time.

### Dramatic shift in chromatin during the maternal-to-zygotic transition

We used three measures of genome-wide recovery of each histone mark to examine their dynamics: the total normalized number of *D. melanogaster* reads (Figure 2A), the number of regions scored by MACS as enriched (Figure 2B) and the total ChIP signal in all enriched regions (Figure 2C). These all gave qualitatively similar results except for H4K5ac, which had anomalously few peaks at early stages despite being found at uniformly high levels across the genome.

**Figure 2.**
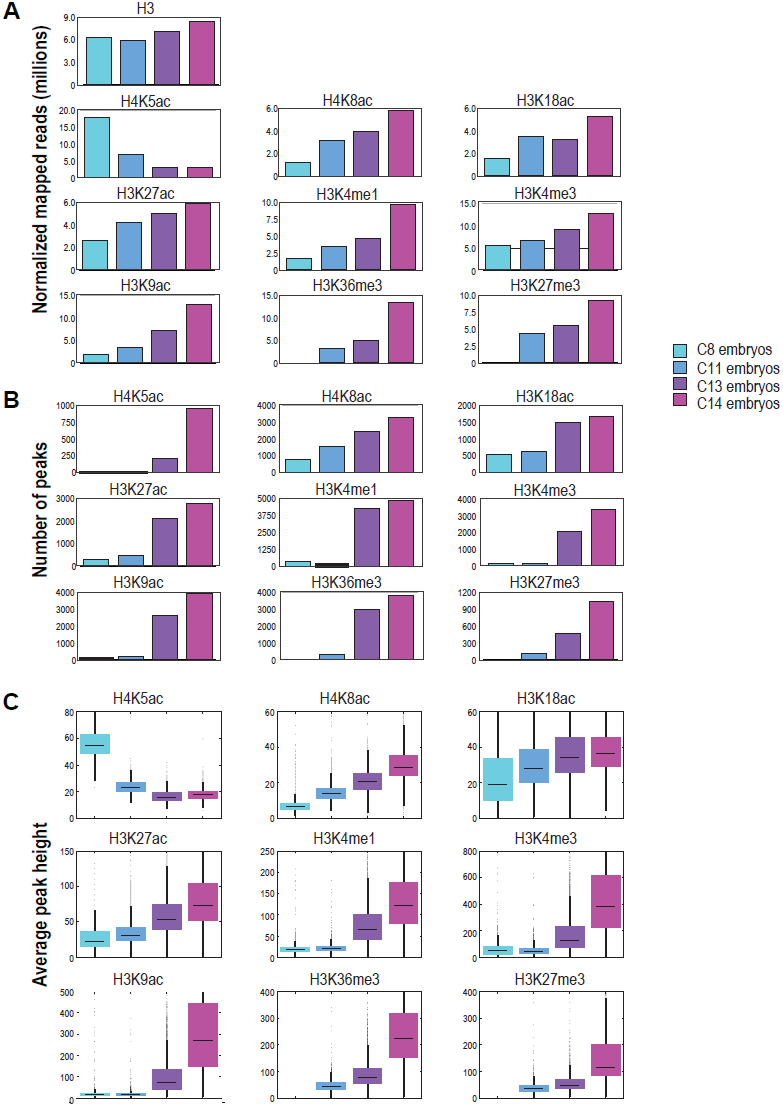
Global levels of histone marks change over early development. (A) The number of aligned reads (after normalization to *D. pseudoobscura*) for the four developmental time points are indicated for each histone mark and histone H3. (B) The number of peaks detected using the peak calling program, MACS (Zhang Y, 2008), for each histone mark at each stage are shown. (C) Box plots show the trend of average ChIP-seq signals over +/-1 kb sequences around the peaks detected across all stages for each histone mark. The dark line in the middle of the plot represents the median, the edges of the box represent the 1st and 3^rd^ quartiles.

As expected, global levels of histone H3 were relatively stable, although we observed a gradual increase of approximately 1.4 fold over time, possibly reflecting an overall increase in compaction of chromatin in cycle 14 relative to cycle 8. The replication associated mark H4K5ac (Sobel et al., 1994), found ubiquitously across the genome, declined rapidly from cycle 8 onwards, consistent with the elongation of cell cycles over time and the decreasing fraction of nuclei caught in S phase. The remaining marks all showed dramatic increases over the MZT. H4K8ac, H3K18ac, and H3K27ac were enriched at hundreds of loci at cycle 8 and steadily increased through cycle 14. The remaining marks, H3K9ac, H3K3me1, H3K4me3, H3K36me3 and H3K27me3, were effectively absent at cycles 8 and 11, but showed sharp increases at cycle 13. This distinction between these two groups of marks is evident when examining levels of histone modification at individual loci (Figure 3).

**Figure 3.**
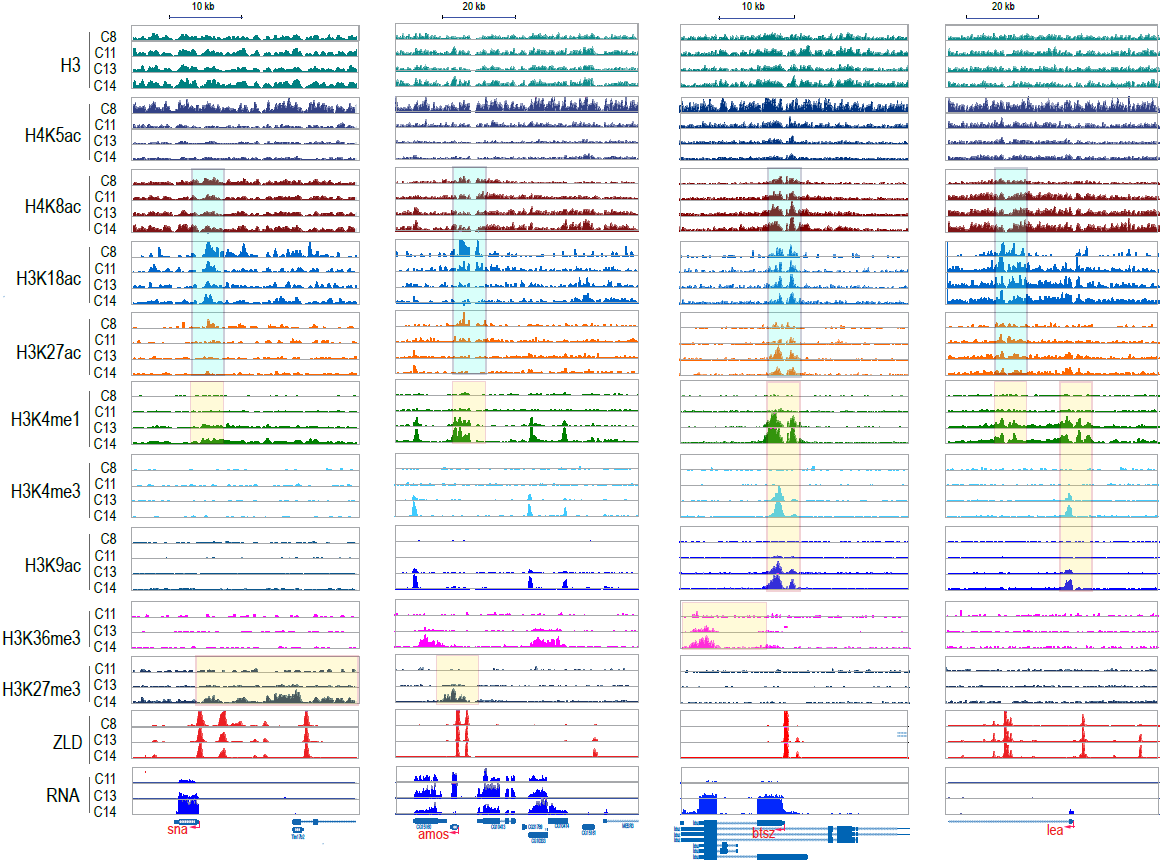
Dynamics of H3 and histone marks around selected genes. The normalized ChIP-seq signal profiles, for histone H3 and nine different histone marks at four development time points at selected genomic loci. Shown are the early onset genes, sna and amos, and late onset genes, btsz and lea. The peak regions of histone acetylation marks detectable prior to MBT are highlighted with cyan-colored boxes. The peak regions for histone marks detected only after the MZT are highlighted by yellow-colored boxes. Below are the ZLD ChIP_seq profile (Harrison et al., 2011) from c8, 13, and 14 embryos, as well as RNA_seq signals (Lott et al., 2011) at c11, c13, c14(b).

### Chromatin changes in transcribed regions are associated with gene activation

The transcription of several thousand genes is initiated during the period covered by our analyses, and we were interested in the relationship between the timing of the onset of transcription at individual loci and their chromatin dynamics. We used high-temporal resolution expression data previously collected by our lab (Lott et al., 2011) to divide the genome into six functional groups of genes - four temporal groups of zygotic genes according to their onset times, as well as maternally deposited transcripts and genes with no detectable transcription levels (Figure 4).

**Figure 4.**
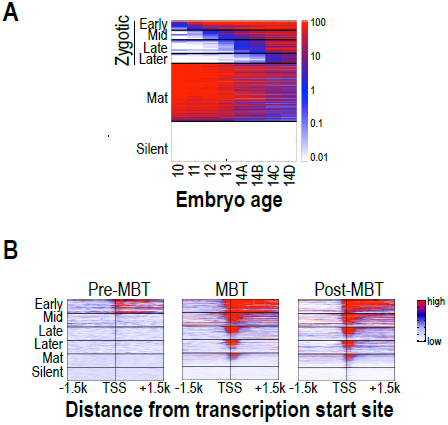
Classification of genes based on timing of transcriptional initiation during early embryogenesis. (A) Gene grouping based on RNA-seq profiles. Based on previous single-embryo RNA-seq data (Lott et al., 2011), we identified three classes of genes: maternally deposited (maternal), zygotically transcribed but not maternal (zygotic) and not expressed in embryo (silent). For the purpose of this study, we further divided the zygotic group into early, mid, late, and later subgroups based on the first mitotic cycle in which transcript is identified. (A) Heatmap showing normalized expression (Lott et al., 2011) of genes in each class at from mitotic cycle 10 through 14. (B) RNA polymerase II ChIP-seq signals (Chen et al., 2013) around the transcription start sites (+/-1.5 kb) of the genes in each category for three developmental time points, pre-MBT (left), MBT(middle), and post-MBT(right) embryos. The genes in each group were ordered based on cycle 14 RNA polymerase II signals (genes with the highest signal are on top).

For genes in each of these different classes, we examined patterns of nucleosome enrichment and histone modifications around the transcription start sites, and in the gene body (Figure 5). Nucleosome free regions (NFRs; areas of relatively low histone H3 recovery) emerged around the transcription start sites of zygotically transcribed genes at roughly the same time that their transcripts were evident in our transcription data (Figure 5). Several histone modifications also appeared along with transcription: H4K8ac, H3K18ac and H3K27ac (Figure 5). In contrast, H3K9ac and the four histone methylation marks examined here were absent at all six classes of genes until cycle 13, increasing substantially in their density as widespread transcription begins during cycle 14.

**Figure 5.**
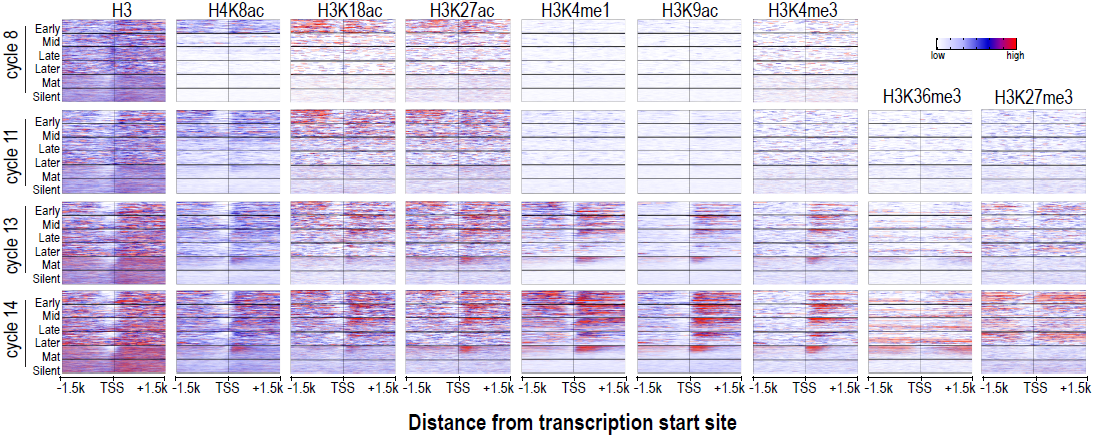
Relationship between H3 depletion, histone modifications and transcription dynamics. Heatmaps show ChIP-seq signals for histone H3 and different histone modification marks at each stage centered around the transcription start sites (+/-1.5 kb). Genes are divided based on their timing of expression as described in Figure 4. For each histone mark and for histone H3 the same scale was used for heatmaps across all four developmental time points. The graphing height and gene order are the same as in Figure 4. TSS = transcriptional start site.

Given the high-resolution of our transcription data (Lott et al., 2011) coupled with our stage-specific identification of histone modifications, we can relate transcriptional activity with the presence or absence of specific histone marks. Despite the generally assumed correlation between transcriptional activation and the presence of H3K4me3 at promoters and H3K36me3 in gene bodies, our data demonstrate that at early expressed genes transcription proceeds in the absence of these marks (Figure 5). It is not until cycle 13 or 14 that H3K4me3 and H3K36m3, respectively, were found associated with actively transcribed genes (Figure 5).

Surprisingly, we observed NFRs at the promoters of maternally deposited genes at all time points (Figure 5), even though there is no evidence that genes in this set are transcribed either from expression data or ChIP with RNA polymerase (Figure 4, Lott 2011, Chen, 2011). The transcription independent nucleosome depletion at the promoter is unique to maternal genes and not seen for the silent gene group. Although maternal genes have NFRs at early time points, the acetyl marks that are associated with the appearance of NFRs in early zygotic genes only appeared at maternal genes at cycle 13, along with RNA polymerase. Based on these observations, the NFRs associated with the promoters of the maternal genes in early embryos may be due to the intrinsic properties of the DNA sequences rather than transcription factor binding.

In the cellular blastoderm, regions both upstream and downstream of the promoters of zygotically expressed genes were enriched for the polycomb-associated mark H3K27me3 (Figure 5). In contrast, H3K27me3 was almost completely absent from maternal genes. The existence of this modification associated with repressed transcription in a transcriptionally active region has been dubbed the “balanced” state (Schwartz et al., 2010), even though it is not clear to what extent such “apparent” coexistence is due to heterogeneity of the cell population present in blastoderm embryos. The function of this “balanced” state is not clear. It is conceivable that in blastoderm embryos this may represent an intermediate step toward full silencing of the genes. In addition, we have found that many genes in the “Later” gene group are expressed only at low levels and are associated with paused RNA polymerase. In these cases, the H3K27me3 may be important for RNA polymerase pausing as suggested by a recent study (Chopra et al., 2011), which poise the gene for “full” activation in later development stages.

### Dynamic histone marks define blastoderm enhancers

Many of the genes transcribed by cycle 14 are expressed in clear spatial patterns (Combs and Eisen, 2013; Lécuyer et al., 2007; Tomancak et al., 2007) driven by the action of distinct transcriptional enhancers. Although several catalogs of blastoderm enhancers exist (Gallo et al., 2011), they are limited in scope. To generate a larger set of likely enhancers, we took advantage of the strong correlation between the binding of transcription factors known to regulate blastoderm expression and enhancer activity (Fisher et al., 2012; MacArthur et al., 2009). For this, we calculated the cumulative in vivo binding landscape of 16 early developmental transcription factors, including the anteroposterior regulators Bicoid, Caudal, Hunchback, Giant, Krüppel, Knirps, Huckebein, Tailless, and Dichaete; and the dorsoventral regulators Dorsal, Snail, Twist, Daughterless, Mothers against dpp, Medea, and Schnurri (MacArthur et al., 2009). We then identified a set of 784 regions showing the strongest overall binding, excluded peaks overlapping promoters and coding regions, and obtained a stringent set of 588 intergenic and intronic putative enhancers.

During cycle 14, when these enhancers are clearly bound by multiple transcription factors and are active, they showed a region of strong nucleosome depletion corresponding to the binding sites for transcription factors. At this stage, the flanking nucleosomes were enriched with characteristic enhancer marks H3K4me1 and H3K27ac as well H3K18ac (Figure 6). These putative enhancer regions frequently occurred within larger regions containing high levels of the repressive mark H3K27me3, and were associated with low levels of H3K4me3 and H3K36me3 (Figure 6).

**Figure 6.**
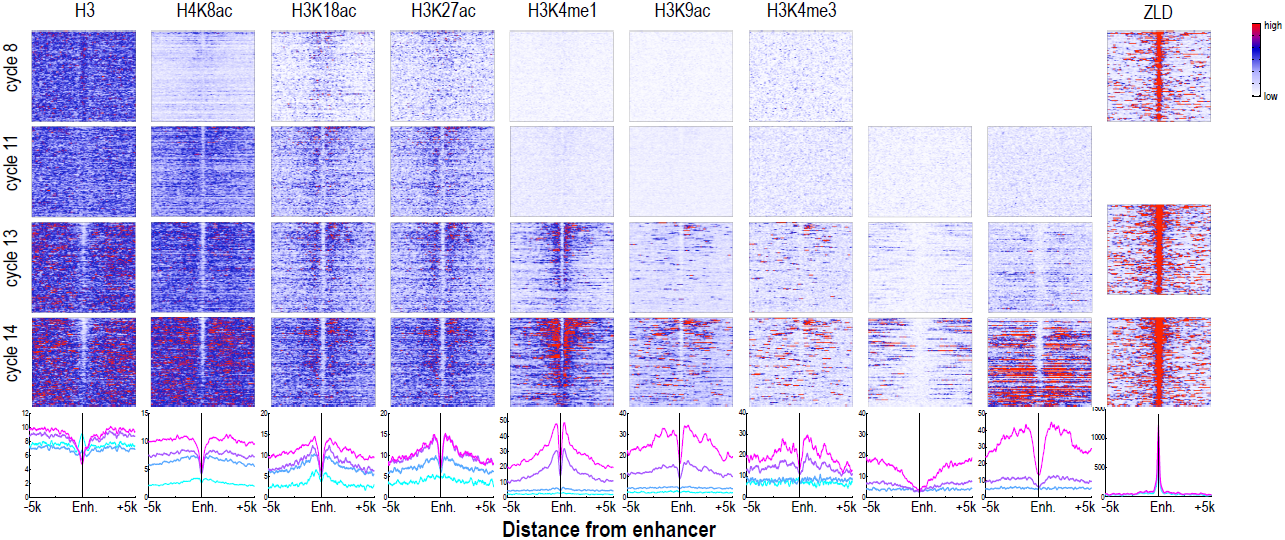
Dynamics of histone H3 depletion and histone modifications around blastoderm embryo enhancers. Heatmaps show ChIP-seq signals for histone H3 and different histone modification marks at each stage centered around putative enhancers (as described in text). Enhancers are ordered by chromatin accessibility, as measured by DNaseI–seq signals from cycle 14 embryos (Thomas et al., 2011) from high (top) to low (bottom). On the right, the heatmaps show the ChIP_seq signals for ZLD around these enhancers at c8, c13, and c14 (Harrison et al., 2011). The line plots at the bottom show the average ChIP-seq for histone H3, histone modifications, and ZLD at each stage around the enhancers. Enh = enhancer.

These identified enhancers, which are destined to be strongly nucleosome depleted at cycle 14, generally had relatively high nucleosome densities at mitotic cycle 8 (Figure 6; top left). At this early time point, flanking nucleosomes were weakly enriched for the three early appearing histone acetyl marks, especially H3K18ac, with these marks becoming more strongly enriched by cycle 11. The process of nucleosome depletion was initially evident at cycle 11, but was much stronger at cycle 13, when these enhancers begin to be active. The enhancer-associated mark H3K4me1 appeared on flanking nucleosomes by cycle 13, but the repressive H3K27me3 did not appear in surrounding regions until cycle 14. This raises the possibility that early events, reflected by the appearance of these enhancer-associated acetylation and methylation marks, play an important role in keeping these regions active once broader domains of inactivity are established.

### Early appearing chromatin features are associated with binding sites for the transcription factor ZLD

As expected from the previous study (Harrison et al., 2011), the set of enhancer sequences described above are strongly associated with early binding of the transcription factor ZLD (Figure 6). This suggests that at least at the enhancers, ZLD binding is likely to play a major role in directing the deposition of the histone acetylation marks in early embryos. To investigate this further and to identify other factors that may play a major role in determining overall histone acetylation patterns, not just the enhancers in early embryos, we carried out kmer enrichment analysis and used the motif search tool MEME to identify sequence motifs associated with different histone mark peaks identified at each stage. We found that the motif most strongly correlated with the early appearing marks, H3K27ac, H3K18ac, H4K8ac and H3K4me1, was ZLD’s CAGGTAG binding site (Figure 7A). A small number of other motifs also show modest enrichment using these two methods, but they failed to show substantial enrichment when the enrichment is plotted around the histone mark peaks. These analyses thus suggested a close connection between ZLD binding and early histone acetylation in general, which is further highlighted by the extremely high degree of overlap between early ZLD-bound peaks and early (cycles 8 or 11) peaks for H3K27ac, H3K18ac and H4K8ac (Figure 7B). The relationship is quantitative, with higher levels of ZLD binding coupled to increased levels of the same three marks in cycle 8 and cycle 11 embryos(Figure 7C). The relationship between ZLD binding and these histone marks decays over time (Figure 7D), likely reflecting the increasingly complex transcriptional profile of the genome. However, the strength of this association in early stages of the MZT suggests that ZLD is a – if not the - dominant factor shaping the early chromatin landscape.

**Figure 7.**
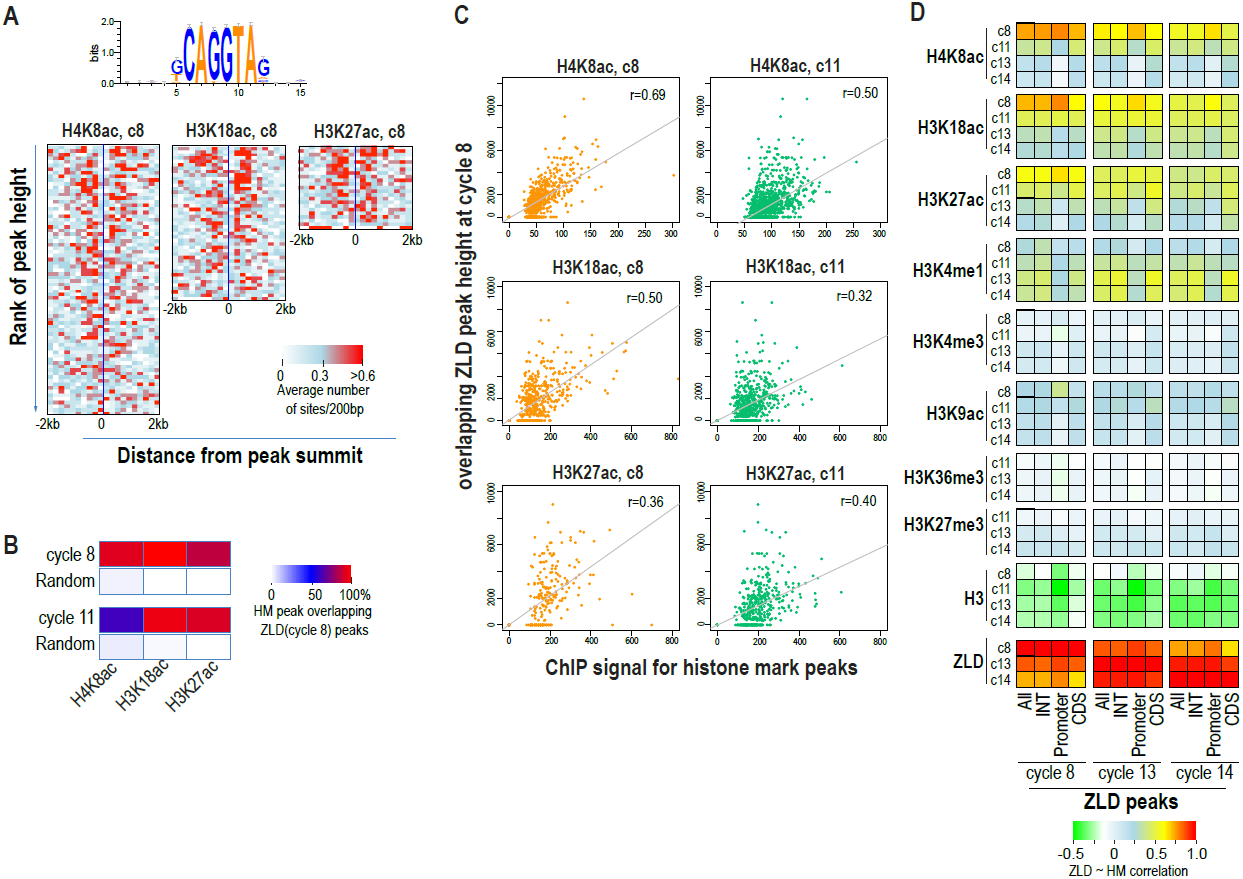
Relationship between histone occupancy, histone modification pattern and ZLD binding. (A) ZLD DNA binding motif enrichment around cycle 8 peaks for H4K8ac and H3K18ac, H3K27ac. Peaks were ranked based on peak height, divided into bins of 100, and the average of the numbers of ZLD binding motif at each position around the peaks in each bin plotted. (B) Heatmap showing the overlap between histone acetylation peaks detected at cycle 8 and 11, and ZLD peaks detected at cycle 8 (top 2000 ranked peaks). As a control, overlaps between histone mark peaks with random set of genomic positions that matched the number of ZLD peaks are shown. (C) Scatter plots showing the correlation between the signals around the peaks for the histone acetylation marks at cycle 8 and cycle 11 and the peak heights of the associated ZLD peaks (within 1 kb of the histone mark peaks). The signal for each peak was the average over the +/-1 kb region surrounding the peak. The correlation coefficient (r) for each plot is shown. (D) Heatmaps shows the correlation between the height of the top 5000 non-redundant ZLD peaks and ChIP-seq signals for histone H3 (averaged over a +/-200 bp region around each ZLD peak) or for each of the histone marks (average signals over +/-1 kb around the ZLD peaks). The correlation coefficients were determined individually for all the ZLD peaks (All), for ZLD peaks that are located in intergenic and intronic (INT) regions, for the promoter (-300 -+100 bp around transcription start site), or for ZLD peaks within coding sequences (CDS).

Beginning at cycle 13, ZLD peaks are generally associated with H3K4me1, which marks enhancers as well as promoters. Thus ZLD binding to enhancers and promoters occurs prior to the deposition of this histone modification. There was also a striking relationship between early ZLD binding and the later presence of the Polycomb-repression associated mark H3K27me3. At cycle 14, a large fraction of H3K27me3 domains overlap with strong ZLD peaks (Figure 8), both at promoters and in intergenic regions. Within the broad H3K27me3 domains, the signal was almost completely absent in regions corresponding to ZLD-binding peaks. This local lack of H3K27me3 in large H3K27me3 regions may reflect an intermediate step toward silencing of the ZLD-associated enhancers that are active in pre-cellular blastoderm embryos, such as the *tll* K10 enhancer (Fig 8C). In other cases where the genes will become transcriptionally active only in later stages, the prevention of local H3K27me3 deposition by ZLD at the putative enhancers marked by ZLD binding may be important to poise them for later activation.

**Figure 8.**
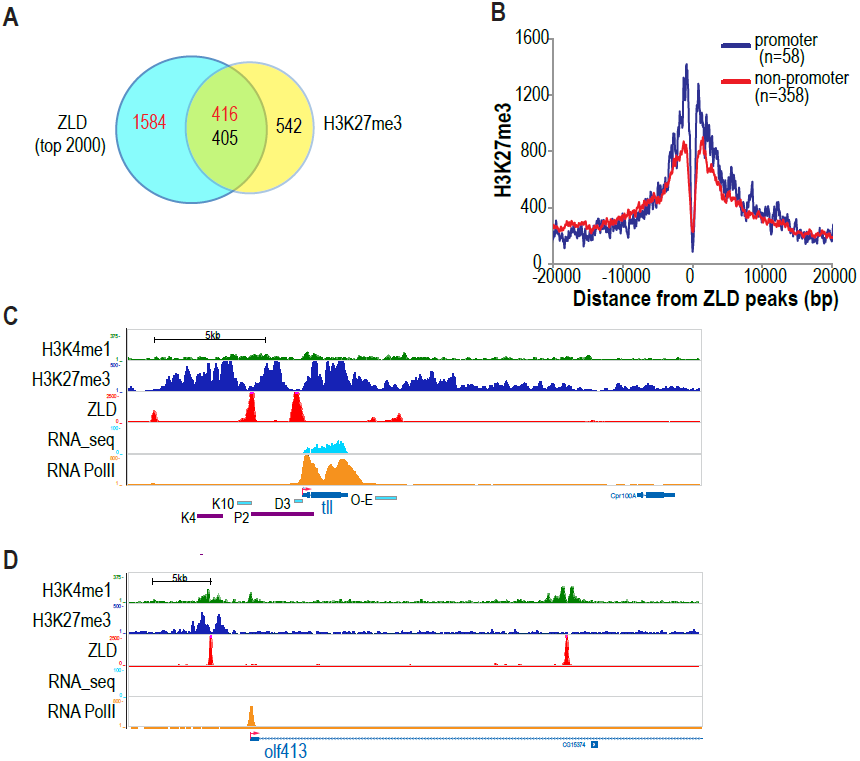
Effect of ZLD binding on H3K27me3 deposition. (A) Venn diagram showing the overlap between H3K27me3 peaks and strong ZLD-binding peaks (top 2000 by rank) at cycle 14. Red numbers indicate ZLD peaks, and black numbers indicate H3K27me3 peaks. (B) Average H3K27me3 ChIP-seq signal profiles around ZLD peaks (promoter peaks shown in blue and non-promoter peaks in red) showing that ZLD peaks tend to occur in large H3K27me3 domains. (C) ChIP-seq profiles at cycle 14 for H3K27me3, ZLD, H3K4me1 and RNA polymerase II along with RNA-seq signals around the gene *tll*. *tll* is expressed both in blastoderm stage embryos and in neuronal cells in later development. Enhancer sequences driving *tll* expression in either the blastoderm (cyan; K10, D3) or in older embryos (purple; K4, P2) are shown (Rudolph et al., 1997). (D) The region centered around the 5’ region of the gene *olf413* is depicted as in (C). At cycle 14, RNA polyermase is paused at the transcription start site of *olf413,* and *olf413* is not transcribed at this time. *olf413* has two putative promoter-distal enhancers that are marked by H3K4me1 and ZLD binding, with one enhancer also associated with a strong H3K27me3 signal.

One good example for such relationship between ZLD binding in a H3K27m3 domain in enhancer posing is shown by the *olf413* gene (Fig.8D). This gene is associated with paused RNA polymerase in blastoderm embryos, and is transcribed only after stage 11 (BDGP). It has two putative enhancers as indicated by the presence of two promoter distal H3K4me1 peaks. One of these peaks also overlapped with H3K27me3 binding, similar to other previously described typical poised enhancers (Calo and Wysocka, 2013). Importantly, in the H3K27me3 region, there is also a strong ZLD binding peak, and around this peak H3K27me3 signal is locally depressed. The potential role of ZLD in poising this enhancer for later activation can be illustrated by the following scenario: conceivably, in later developmental stages, in the cells where ZLD continues to be expressed, ZLD binding will continue to maintain the enhancer in a poised state or lead to activation of this enhancer. In contrast, in cells that do not express ZLD, this gap of H3K27me3 will be filled leading to complete silencing of this enhancer.

### Histone marks at enhancers are decreased in embryos lacking maternal ZLD

To directly analyze the role that ZLD plays in the activation of the zygotic genome during the MZT, we carried out a limited series of ChIP experiments using embryos lacking maternal *zld* RNA, obtained from *zld*-germ-line clones (Liang et al., 2008). These females lay significantly less than their wild-type counterparts, and thus obtaining sufficient amounts of staged chromatin was a challenge.

ChIP with an anti-ZLD antibody on these nominally *zld-* embryos at mitotic cycle 12 to mid-14, showed modest ZLD binding at the same set of sites bound in wild-type embryos (Figure 9). This residual ZLD activity is likely due to zygotic transcription of the paternal copy in female embryos. (*zelda* is on the X chromosome and thus male offspring do not receive a functional *zelda* from their father.) Thus these ChIP data reflect the depletion, rather than elimination, of ZLD.

**Figure 9.**
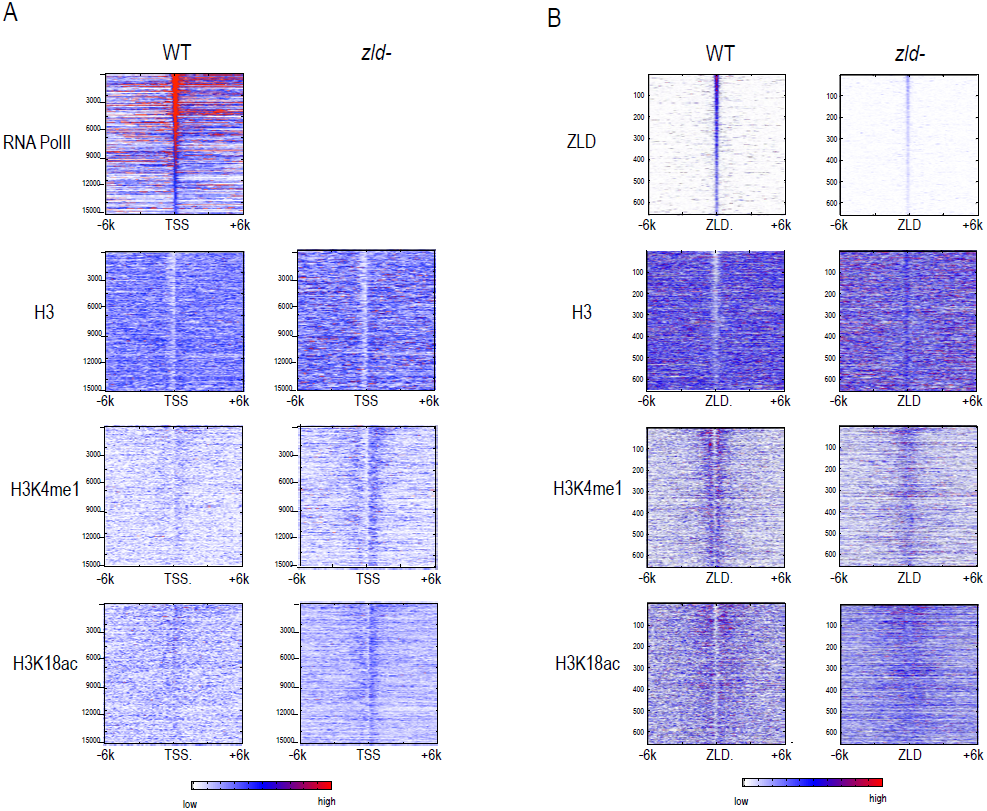
Effect of zld mutation on histone occupancy and modifications. Heatmaps show ChIP-seq data from WT embryos (left) and embryos lacking maternal *zld* (right). (A) Heatmaps centered at transcription start sites ordered by cycle 14 RNA polymerase II binding. (B) Heatmaps centered around ZLD (Harrison et al., 2011) for the top 1000 bound regions and are centered around the intergenic ZLD binding site. TSS, transcription start site

Intergenic ZLD-bound regions (Figure 9B) had a marked loss of H3K4me1 and a decrease of H3K18ac. The ZLD-associated NFR present in wild-type embryos was almost completely gone. In contrast, we saw a limited effect of ZLD depletion on the promoter histone state globally. There is still a strong NFR, and the marks we observed are present at roughly the same levels.

## Discussion

### Nature of chromatin changes during MZT

During the first stage of embryonic development, the genome must be reprogrammed from the differentiated states associated with the egg or sperm to create a set of totipotent cells capable of generating a new organism. During this early phase of development, gene expression is limited. It is currently unknown how the regulation of gene expression relates to the reprogramming of the epigenetic state and what factors might regulate these changes in chromatin architecture. By combining high-resolution gene expression analysis with precise mapping of nine histones mark through this early stage of development, our data suggest that this reprogramming, at least in *Drosophila*, occurs by transitioning through a naïve state in which many histone marks commonly present in somatic cells are absent or at comparably low levels. We further demonstrate that histone acetylation of H3K18, H3K27, and H4K8 precedes most histone methylation. Thus we suggest that the establishment of the totipotent chromatin architecture proceeds in an ordered process with acetyl marks being deposited prior to methyl marks.

Studies in other organism have similarly suggested that this early reprogramming is characterized by a transition through a relatively unmodified chromatin state. Immunostaining in mouse and bovine embryos has demonstrated that some histone methylation marks are removed following fertilization (Burton and Torres-Padilla, 2010). Additionally, studies in zebrafish have demonstrated widespread changes in chromatin marks as the embryo progresses through the MZT. While the extent and location of specific histone modifications in zebrafish is not consistent between recent studies (Lindeman et al., 2011; Vastenhouw et al., 2010), a general widespread increase in histone methylation (H3K4me3, H3K27me3, H3K36me3, and H3K9me3) is evident at the MZT. Thus in most, if not all, organisms studied to date there is a dramatic increase in the abundance of histone modifications at the MZT, coinciding with zygotic genome activation.

One important lingering question is how this naïve state is established; whether there is a specific system that removes gametic marks associated with sperm and egg at fertilization, or whether the rapid replication cycles of early development are simply incompatible with active and differentiated chromatin. The lack of some histone marks in developing mouse and bovine embryos, which do not undergo rapid cell cycles early in development, suggests that while the mechanisms may vary between species the removal of parental histone modifications may be a general feature of reprogramming. In the future, it will be important to understand how this transition is regulated to allow for the generation of a totipotent cell population.

The data presented here provide only a glimpse into this process, but the biochemical and genomic tools exist now to create a more complete accounting of the proteins that interact with DNA during early embryogenesis, to understand how the process is regulated, and to determine the effect this process has on subsequent steps in development.

### Early transcription does not require marks canonically associated with activation

Post translational modifications on histones have been associated with differential levels of gene expression and have been used to predict gene activity. H3K4me3 at promoters and H3K36me3 within gene bodies widely reflect active transcription. However, our data demonstrate that transcription can occur in the near absence of these marks. Through the combination our previously published high-resolution gene expression data and our precise mapping of histone modifications, we show that at the promoters and within the gene bodies of those genes transcribed prior to cycle 14 that H3K4me3 and H3K36me3 levels are below detection. Interestingly global H3K36me3 levels are also below the level of detection in the mouse embryo as it undergoes the first wave of zygotic genome activation (Boskovic et al., 2012).

Together these data suggest that marks canonically associated with gene activation at later stages of development are not associated with transcriptional activity in the very early zygote. Moreover, recent data has demonstrated that these marks may not be required for transcription later in development. For example, in the Drosophila wing disc methylation of H3K4me3 is not required for transcriptional activity (Hodl and Basler, 2012). Additionally, data from *S. cerevisiae* show robust transcriptional activation of a gene localized to heterochromatin in the presence of minimal amounts of H3K36me3 (Zhang et al., 2014). Thus increasing evidence suggests that while specific histone modifications may be associated with levels of gene activity they are not required, in at least some cell types, for expression.

By contrast, in zebrafish H3K4me3 was identified at promoters of genes prior to their occupancy by RNA polymerase (Lindeman et al., 2011; Vastenhouw et al., 2010). Because those genes marked by H3K4me3 early are more likely than most genes to be activated at the MZT, it has been proposed that this mark is preparing genes for activation in the early zebrafish embryo. Thus, while histone marks can be associated with specific transcriptional outputs it appears that they are neither necessary nor sufficient for predicting gene expression.

### A simple model for early genome activation

While the data presented here are far from complete, together with our previously published high-resolution transcriptional analysis, they suggest a model in which regions of genomic activity in the cellular blastoderm are established by events that transpire earlier in development. In particular, they indicate that the early binding of ZLD to target sites across the genome may trigger a cascade of events – reflected in the early histone depletion and the appearance of several histone acetylation marks, and the subsequent appearance of functional class-specific methylation marks – that may act to counter the establishment of Polycomb-mediated repression in many loci (Figure 10).

**Figure 10.**
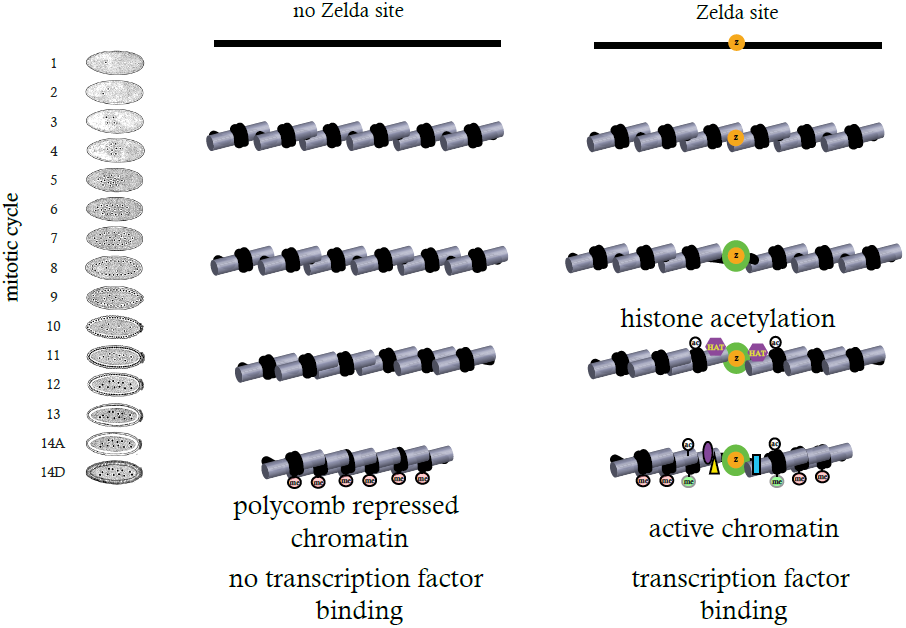
Model for ZLD function during zygotic genome activation. ZLD binds to enhancers in pre-MBT embryos at as early as cycle 8. This leads to histone acetylation and nucleosome remodeling around ZLD binding sites, which facilitates binding by other transcription factors, and in many other cases leads to additional deposition of histone marks including H3K4me1 while at the same time prevents local deposition of repressive histone mark H3K27me3 and presumably formation of repressive higher order chromatin structure.

There are obviously other players in the process besides ZLD, as evidenced by the maintenance of many features of an active state at promoters in ZLD depleted embryos. However, the tight coupling of ZLD binding and early histone acetylation suggests that ZLD is playing a major role in the process.

Of course our data are largely silent as to how ZLD might be playing this role. That the first feature we observe associated with ZLD binding is histone acetylation, raises the possibility that ZLD specifically recruits histone acetyltransferases, several of which including Nejire and Diskette are maternally deposited, and that these modifications play a direct or indirect role in genome activation. Alternatively, ZLD may simply act as a kind of steric impediment to subsequent chromatin compaction and silencing – with the observed histone acetylation an indirect byproduct of early ZLD binding.

We have previously observed that, while ZLD binding is fairly stable across the MZT, some of the regions it binds at cycle 8 are unbound at cycle 14 (Harrison et al., 2011). This may reflect the need for other factors to work in conjunction with ZLD while more restrictive chromatin is established. Indeed, while ZLD protein levels remain high through the MZT, the increasing number of nuclei means that absolute ZLD levels are dropping in each nucleus and may reach a point at which ZLD binding alone is insufficient to keep regions active or resist silencing.

The possible interaction between ZLD and Polycomb repression may help explain some puzzling and inconsistent results we and others have obtained about the effects of ZLD on expression driven by transgenic enhancer constructs. While the removal of ZLD sites from transgenic enhancers almost always reduces their activity, it almost never eliminates it, which would seem to invalidate the model we have presented above. However, the transgenic enhancer constructs that have been tested in this regard invariably have insulator sequences that resist the spread of Polycomb silencing, and have been inserted into parts of the genome that have been pre-screened to be relatively free of such silencing. If ZLD’s primary role is to resist Polycomb silencing, then we would not expect the loss of ZLD binding sites to affect the activity of enhancers placed into such activation friendly environs.

### ZLD as a pioneer transcription factor

Work from Ken Zaret and others over the past decades has identified a class of transcription factors, known as “pioneer” factors, that bind early to enhancers during differentiation and thereby promote the binding of other factors to the enhancer. Zaret attributes two characteristics to pioneer factors: 1) they bind to DNA prior to activation and prior to the binding of other factors, and 2) they bind their target sites in nucleosomes and in condensed chromatin (Zaret and Carroll, 2011).

ZLD clearly has the first characteristic. But it is not clear that is has the second. Our data suggest that there is essentially no condensed chromatin in the early embryo, as nucleosome density is relatively low, nucleosomes are relatively evenly distributed across the genome, and hallmarks of repressed chromatin are absent. This is consistent with the unusually broad binding of ZLD to its target sequences: ZLD binds to more than fifty percent of its target sites in the genome, far more than what is typical for other factors later in development. We and others have shown that the restricted binding of other factors is largely due to the occlusion of most of their sites by condensed chromatin. Perhaps ZLD binds to a large fraction of its sites because there simply is no condensed chromatin in the early embryo. If so, ZLD would not require, and therefore would likely not possess, the ability to bind its sites in condensed chromatin.

Nonetheless, it is clear, if our model is correct, that ZLD is fulfilling the same general role that pioneer factors carry out – getting to the genome first and facilitating the subsequent binding of other factors.

While there are no clear ZLD homologs outside of insects, it has recently been shown that in zebrafish the transcription factor Pou5f1, in combination with Nanog and SoxB1, drives zygotic genome activation and may share with ZLD a pioneer-like activity (Lee et al., 2013; Leichsenring et al., 2013). Together these data suggest that pioneer transcription factors may generally be required to prepare the embryonic genome for widespread transcriptional activation at the MZT. Interestingly, Pou5f1 is homologous to the canonical pluripotency factor Oct4, which along with Nanog and Sox2, are transcription factors expressed to generate induced pluripotent stem cells. Together these data make an explicit connection between the role of pioneer-like factors in zygotic genome activation and the establishment of a totipotent state.

### Separation of enhancer specification from output

Another attractive feature of the model presented above is that it would explain an important and unexplained question about transcriptional enhancers: given that essentially every enhancer sized stretch of the *Drosophila* genome contains a large number of binding sites for the factors active in the cellular blastoderm (or any other stage of development) (Berman et al., 2002), why is it that only a small fraction of the genome functions as an enhancer?

It has long been thought that the difference between enhancers and the remainder of the genome is that enhancers do not simply contain binding sites, but rather have these sites in a particular configuration that leads to activation. However, the arrangement of binding sites within *Drosophila* enhancers is highly flexible (Hare et al., 2008; Ludwig and Kreitman, 1995; Ludwig et al., 2005; Ludwig et al., 1998), and we have struggled to find any evidence for strong “grammatical” effects in enhancer organization.

The data presented here and elsewhere on ZLD binding and activity support the alternative explanation that the specification of enhancer location and output are distinct processes carried out by specific sets of factors: pioneer factors like ZLD – that determine where an enhancer will be, by influencing the maturation of genomic chromatin, and more classical patterning factors that determine what the transcriptional output of the enhancer will be.

## Materials and Methods

### Antibodies

The antibodies for histone H3, and various histone modifications were purchased from commercial sources as listed in Table 1.

### Fly strains

The zld^294^ mutant and the ovo^D1^ mutant lines used to obtain *zld* maternal mutant embryos using the FLP-DFS technique (Chou and Perrimon, 1996) have been described previously (Liang et al., 2008) and were obtained from the Rushlow lab at NYU.

### In vivo formaldehyde cross-linking of embryos, embryo sorting, and chromatin preparation

*D. melanogaster* flies were maintained in large population cages in an incubator set at standard conditions (25°C). Embryos were collected for 30 minutes, and then allowed to develop for 55, 85, 120 or 160 additional minutes before being harvested and fixed with formaldehyde. The fixed embryos were hand sorted in small batches using an inverted microscope to remove embryos younger or older than the targeted age range based on morphology of the embryos as previous described (Harrison et al., 2011). After sorting, embryos were stored at -80°C. After all collections were completed, the sorted embryos of each stage were pooled, and a sample of each pool were stained with DAPI. The ages of the embryos and their distribution in the two younger embryo pools (c7-9, and c11-12) were determined based on nuclei density of the stained embryos. The ages of embryos between c13 and c14 were harder to distinguish based on DAPI staining alone, and they were determined based more on morphology. 7.5, 0.7, 0.4, and 0.3 g of embryos at four different stages respectively, were used to prepare chromatin for immunoprecipitation following the CsCl_2_ gradient ultracentrifugation protocol as previously described (Harrison et al., 2011).

### ChIP and sequencing

The chromatin obtained was fragmented to sizes ranging from 100 to 300 bp using a Bioruptor (Diagenode, Inc.) for a total of processing time of 140 min (15 s on, 45 s off), with power setting at “H”. Prior to carrying out chromatin immunoprecipitation, we mixed the chromatin from each sample with a roughly equivalent amount of chromatin isolated from stage 4-5 (mitotic cycle 13 – 14C) *D. pseudobscura* embryos, and used about 2 µg of total chromatin (1 µg each of the *D. melanogaster* and *D. pseudobscura* chromatin) for each chromatin immunoprecipitation. The chromatin immunoprecipitation reactions were carried out as described previously (Harrison et al., 2011) with 0.5 ug anti-H4K5ac (Millipore, 07-327), 0.5 ug of anti-H3K4me3 (Abcam, ab8580), 0.5 ug of anti-H3K27ac (Abcam, ab4729), 1 ug of anti-H3 (Abcam, ab1791), 0.75 ug anti-H3K4me1 (Abcam, ab8895), 0.75 ug anti-H4K8ac (Abcam, ab15823), 1.5 ul of anti-H3K9ac (Activemotif, 39138), 0.75 ug anti-H3K18ac (Abcam, ab1191), 3 ug anti-H3K27me3 (Millipore, 07-449), or 0.75 ug anti-H3K36me3 (Abcam, ab9050). The sequencing libraries were prepared from the ChIP and Input DNA samples using the Illumina TruSeq DNA Sample Preparation kit following the manufacturer’s instructions, and DNA was subjected to ultra-high throughput sequencing on a Illumina HiSeq 2000 DNA sequencers

### Mapping sequencing reads to the genome, and peak calling

Sequenced reads were mapped jointly to the April 2006 assembly of the *D. melanogaster* genome [Flybase Release 5] and the November 2004 assembly of the *D. pseudoobscura* genome [Flybase Release 1.0] using Bowtie (Langmead, 2010) with the command-line options “-q -5 5 -3 5 -l 70 -n 2 -a -m 1 —best-strata”, thereby trimming 5 bases from each end of the 100 base single reads, and keeping only tags that mapped uniquely to the genomes with at most two mismatches. Each read was extended to 130 bp based on its orientation to generate the ChIP profiles. We called peaks for each experiment using MACS (Zhang et al., 2008) v1.4.2 with the options “—nomodel —shiftsize = 130”, and used Input as controls.

### Data normalization

The addition of *D. pseudoobscura* chromatin prior to the chromatin immunoprecipitation provided us with a means to normalize the ChIP signals for each histone mark and for H3 between different stages. To normalize, we first determined the scaling factor needed to normalize the number of reads for *D. pseudoobscura* to 10 million, and scaled the signals of *D. melanogaster* ChIP profile in each sample using this factor. We then multiplied the scaled *D. melanogaster* signals by the ratio of *D. pseudoobscura* reads to *D. melanogaster* reads in the Input sample, which represents the relative amounts of chromatin of the *D. melanogaster* and the *D. pseudoobscura* in the starting chromatin samples used for the chromatin immunoprecipitation reactions.

### Overall dynamics of ChIP signals under enrichment peaks

Starting with peaks called by MACS as described above, we identified subpeaks by peaksplitter [http://www.ebi.ac.uk/research/bertone/software], and generated a consolidated list of subpeaks for each histone mark for all stages by joining each group of subpeaks that are within 200bp into a single peak. We calculated the ChIP signal for each subpeak at each stage by summing the ChIP signal around a 500 bp window center around of each peak position in the normalized ChIP profile generated as described above. To show the overall trend of each histone mark, the range of the ChIP signal among all the subpeaks at each stage is shown as box plot.

### Gene classification according to transcription dynamics

Using our previous single-embryo RNA-seq data from Lott et al. (Lott et al., 2011), genes were classified as zygotic or maternal. In addition, we defined another class of non-expressed (showing no transcription from mitotic cycles 10 through the end of cycle14. Finally, we further divided the zygotic genes into four different groups based on their onset of zygotic expression (first time point with FPKM> 1). This includes 107 genes whose onset of expression was around mitotic cycles 10-11 (“Early” group), 99 genes at cycles 12-13 (“Mid”), 143 genes at early cycle 14 (“Late”), and 99 genes during late cycle 14 (“Later”).

### Defining embryo blastoderm enhancers

A set of putative enhancers was defined based on the in vivo binding locations for early transcription, as measured previously by us using ChIP-chip (MacArthur et al., 2009). Here, we summed the raw ChIP-chip signal for 16 factors, including the A/P (Bicoid, Caudal, Hunchback, Giant, Krüppel, Knirps, Huckebein, Tailless, and Dichaete) and D/V (Dorsal, Snail, Twist, Daughterless, Mad, Medea, and Schnurri) regulators. We then identified all regions with cumulative signal over 20. This yielded 784 genomic regions, with an average length of 488 bp. These putative enhancers were then classified based on their position with regard to nearby genes, retaining only a set of 588 intergenic and intronic putative enhancers.

### Analysis of motif enrichment

Two methods were used to investigate the DNA motifs enriched around the peaks identified by MACS. First, 7mers enriched in the 2 kb sequences around the peaks for each experiment were identified by comparing the frequency of each 7mer to the 7mer distribution in randomly selected 2 kb sequences throughout the genome. The selection of the random sequences was restricted to the major chromosome arms excluding the heterochromatic sequences, and the distribution of the number of random sequences were set to match the distribution of peaks among different chromosome arms. The enrichment of the 7mers was ranked based on Z scores. In parallel, the motif enrichment analysis was also carried out using MEME (Bailey and Elkan, 1994) with motif length set at 6–10 and maximum number of motifs to be found at 10. In this case the sequences located in the 250-650 bp surrounding the maximum of the histone mark peak were used, and random sequences selected using the same criteria as the kmer enrichment analysis were used as negative control. The search was limited to the 150 top ranked peaks for each histone mark. After the candidate enriched motif was identified from these two methods, the motifs were used to map the enrichment around all the peaks by patser (Hertz and Stormo, 1999) using a ln(p-value) cutoff of -7.5, and with Alphabet set at “a:t 0.3 c:g 0.2”.

### ChIP-seq in *zld* mutant embryos

To obtain embryos depleted of maternal *zld* RNA, the FLP-DFS technique was used. Briefly, *zld^294^, FRT19A*/*FM7* (Liang et al., 2008) virgin females were crossed with *ovo^D1^,hsFLP112,FRT19A*/Y (Liang et al., 2008) males. The larvae developed from embryos laid by females from these crosses were heatshocked twice, each for 2 hours at 37°C, when they were between 24-48hr, and between 48-72 hr old. Collection of the mutant embryos from the resulting female progeny, as well as the aging and fixation of the embryos was carried out following standard protocol as described above except that the collection period is 3 hours followed by 1 hr aging. The embryos were sorted to remove deformed post cycle 14 embryos. As a control, wild-type embryos were collected, treated in parallel and sorted to remove embryos older than stage 5. The ChIP-seq was carried out with the chromatin from the mutant and wild-type embryos using anti-H3, anti-H3K18ac, anti-H3K4me1, and anti-ZLD antibodies as described above.

### Data availability

We are setting up a site to distribute all of the data collected as part of this project and submitting the sequencing reads to the relevant databases. In the meantime, we are happy to share any of these data described here.

## Acknowledgements

This was funded by an HHMI Investigator award to MBE. TK is a member of the Israeli Center of Excellence (I-CORE) for Gene Regulation in Complex Human Diseases (Israel Science Foundation grant No. 41/11), and the Israeli Center of Excellence (I-CORE) for Chromatin and RNA in Gene Regulation (grant No. 1796/12).

